# Predicting trophic links across freshwater fish food webs from species traits

**DOI:** 10.64898/2026.07.31.742128

**Authors:** Imran Bin Younos

## Abstract

Body size is the principal trait governing trophic interactions in freshwater fish communities, yet the body size data available for most species are a species-level maximum. Whether this global proxy can predict trophic links in systems the model has never seen is untested. We evaluated four models on 37 freshwater fish food webs under leave-one-study-out cross-validation (LOSO-CV), predicting each held-out network from species traits with no observed interactions. Locally measured body mass ratios predicted links accurately (median ROC-AUC 0.973). FishBase traits predicted fish-associated links nearly as well (0.887 against 0.912 on the same networks). Whole-network scores were lower (0.607); the shortfall was confined to pairs between non-fish taxa, which FishBase does not describe. A habitat overlap score from FishBase depth-stratum categories added no signal, and a graph attention network did not outperform the random forest. For held-out systems within the range of the training corpus, fish-associated links are recovered from globally available traits alone. This did not extend to the three out-of-region networks, where prediction dropped to near or below chance. Cross-region transfer will need a larger and more geographically varied set of fish food webs.

## Introduction

Trophic interactions are the backbone of ecosystem function, determining how energy flows through communities, mediating population dynamics, and governing responses to environmental change (Van der Putten et al. 2004; Thompson et al. 2012; Rosenblatt and Schmitz 2014). Yet the vast majority of freshwater ecosystems worldwide lack any empirical food web data, and this gap is largest precisely where aquatic biodiversity is highest: tropical rivers, lakes of southern Asia, and freshwater systems of sub-Saharan Africa (Thompson et al. 2012; Cameron et al. 2019; Poisot et al. 2021). Gut content analysis and stable isotope approaches remain essential for identifying trophic links, but they are time-consuming and hard to apply at large spatial scales (Davis et al. 2012). Trait-based interaction models can predict trophic interactions and generate first approximations of food webs where empirical data are lacking (Caron et al. 2022).

Species functional traits provide a more direct approach. Body size is the dominant trait organising trophic contacts in aquatic communities. Predation is fundamentally an individual-level encounter process, so the probability of a predator consuming a given prey item scales with the size asymmetry between them (Woodward et al. 2005; Brose et al. 2006; Petchey et al. 2008). Feeding guild, trophic level, and habitat position add discriminating power when species span a range of ecological roles (Gravel et al. 2013). Machine learning models trained on these traits have achieved remarkable accuracy at predicting trophic links. Pomeranz et al. (2019) inferred predator-prey interactions across 17 New Zealand stream food webs using locally measured body size distributions, producing predicted food web structures closely matching their empirical counterparts. Caron et al. (2022) built a trait-based model of the entire European vertebrate food web and found that models calibrated with as few as 100 interactions estimated the full metaweb reasonably well (AUC ≈ 0.92). Van Kleunen et al. (2026) extended this to 290 food webs spanning five ecosystem types, with stacked models combining trait-based and structural predictors achieving near-perfect performance under within-network cross-validation.

These results confirm that trophic links are predictable from traits, but only when one condition holds that is rarely stated. The model already knows which species are present in the target community. In within-network cross-validation, training and test edges are drawn from the same network, so the model has implicitly learned the local species pool, body size range, and trophic diversity of the target system. Regional metaweb approaches share species identities and evolutionary history across training and test data. Both approaches make the problem considerably easier than the real challenge. That challenge is predicting interactions in a community the model has never encountered, using trait information that may not discriminate among the target species. The quantitative consequences of this distinction for freshwater fish food webs have not been established.

Globally accessible databases such as FishBase (Froese and Pauly 2026) provide trait records for most described fresh-water fish species, making them the natural starting point for cross-system prediction. FishBase maximum body length (Lmax) is available for virtually all species and is the most widely used body size proxy in comparative food web research (Brose et al. 2006; Petchey et al. 2008). But Lmax records the largest individual ever documented for a species across its full geographic range and sampling history. It carries no information about the body size structure of any particular population at any specific time, the life-stage composition of the community, or the actual size asymmetry between consumer and prey present in a given lake or stream (Poisot et al. 2021). Whether this global proxy retains enough signal to support cross-system link prediction has not been tested against a locally measured alternative under a controlled, identical evaluation design.

The same problem applies to habitat information. FishBase depth-stratum categories (DemersPelag) summarise where a species has been recorded across its entire range, not where individuals forage in a specific water body. Global trait databases compile species-level central tendencies from occurrence records aggregated across ranges; trophic contact depends on which individuals are present in a specific community at a specific time, a question that species-level global averages cannot reliably answer (Caron et al. 2024). Whether these global habitat descriptors contribute any predictive value for local trophic interactions has not been evaluated in a cross-system framework.

A parallel question concerns whether structural information from training food webs can compensate for trait inadequacy in ecologically novel systems. Graph neural networks trained on food web topology encode regularities about how trophic hierarchy, predator-prey asymmetry, and connectivity are organised across systems (Biton et al. 2025; Strydom et al. 2023). These structural priors could in principle anchor predictions when global trait features fail to discriminate between species. Whether this compensation operates in a fully inductive setting, where no interactions from the target food web are available at any stage, remains untested.

We address these questions using 37 freshwater fish food webs from two independently compiled databases, evaluating four models under leave-one-study-out cross-validation (LOSO-CV). In each fold, every model is trained exclusively on known food webs and applied to a held-out network with zero observed interactions. The design isolates three specific questions. First, how large is the prediction gap between locally measured body mass ratios and FishBase maximum body length as a global proxy, when algorithm and evaluation design are held constant? Second, do vertical habitat overlap scores derived from FishBase depth-stratum categories add predictive value beyond body size and trophic level features, and what does their performance reveal about the structural limitations of global trait databases? Third, does training a graph attention network (GAT) on food web structure improve cross-system link predictions when no structural information is available at test time?

## Methods

### 2.1 Food web data sources

We assembled freshwater food web data from two publicly available databases. The Global Archive of Trait, Evolution, and Energy Use (GATEWAy; Brose et al. 2019) is a curated compilation of 290 published food webs in which every consumer-resource interaction has been verified against primary literature and each node is associated with standardised body mass, movement type, and metabolic type. We downloaded the complete dataset from the GATEWAy project repository (https://github.com/globalbioticinteractions/brose-gateway) and filtered to freshwater systems by retaining all networks designated as aquatic, non-marine habitats. The second source was Mangal (mangal.io; Poisot et al. 2016), an open repository of ecologist-contributed food web datasets. We queried all networks flagged as freshwater using the rmangal R package (3.0.0) and retrieved node and edge records for each.

Before combining the two sources, we removed 18 Mangal networks in which at least one fish node was identified by FishBase as a strictly marine species (freshwaterflag = 0). We then applied a fish-richness criterion to both sources, retaining only those networks containing at least two fish species with a non-missing maximum body length (Lmax) value in FishBase. Networks without this minimum trait coverage produce constant pairwise feature vectors and contribute no information to either training or evaluation, so their exclusion improves data quality without biasing the scope of inference. After both filtering steps and removing duplicate networks that appeared in both databases, the combined corpus comprised 37 networks: 34 from GATEWAy and 3 from Mangal, encompassing 2,155 nodes and 22,373 consumer-resource interactions across 50 unique fish species (Figure 1).

**Figure 1.**
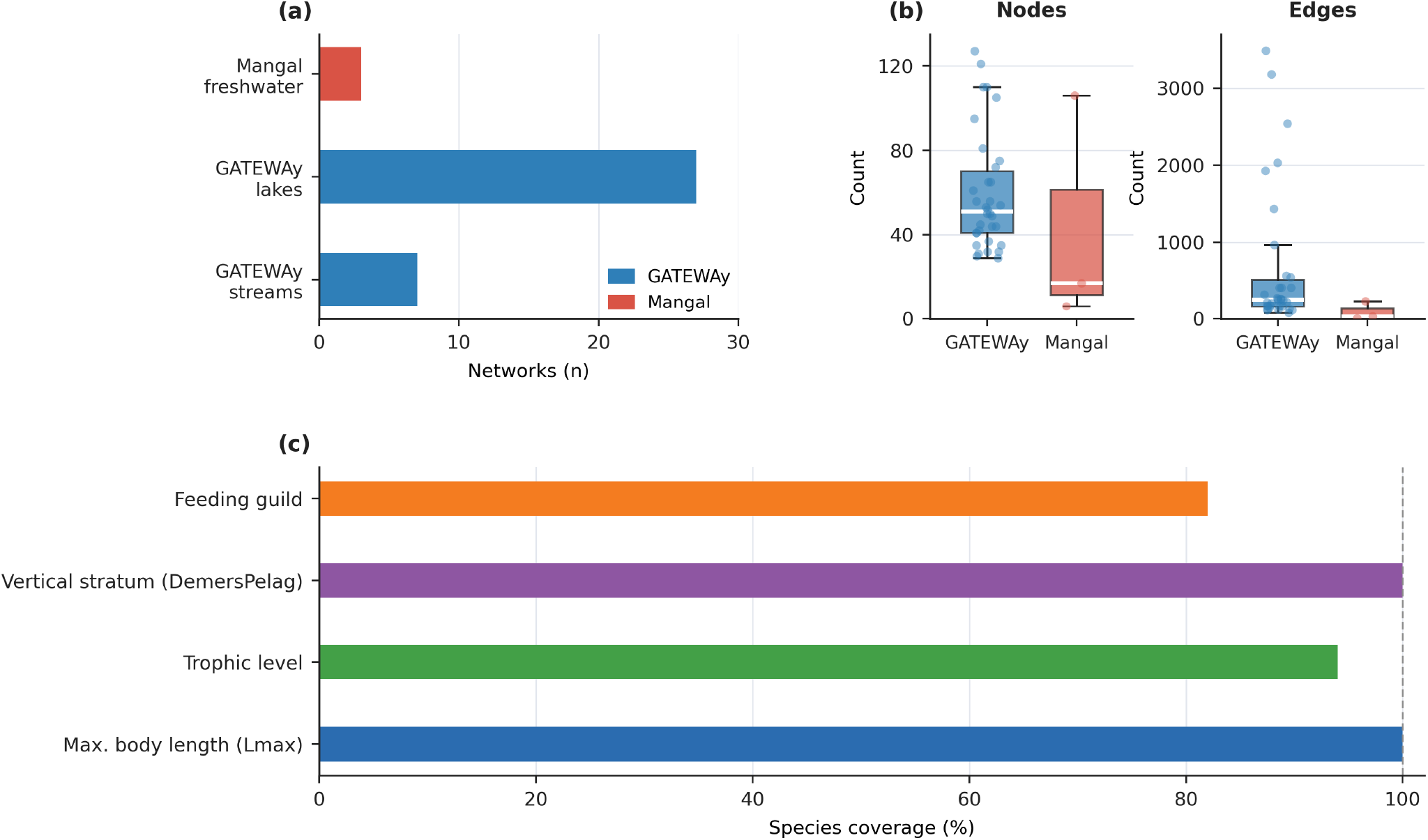
Characteristics of the 37-network freshwater fish food web corpus. (a) Number of networks by source database and ecosystem type; each bar represents a single source-ecosystem combination. (b) Distribution of node and edge counts per network. Boxes show interquartile range; white bars show medians; points are individual networks (jittered). (c) FishBase trait coverage across the 50 fish species included in the corpus.

### 2.2 Fish species trait data

Fish species trait data were retrieved from FishBase (Froese and Pauly 2026; accessed June 2026) using the rfishbase package (5.0; Boettiger et al. 2012) in R (4.3.1). For every fish species identified across the 37 networks, we retrieved maximum total body length (Lmax, cm; sourced from the Length table with length type LTypeMaxM as the primary measure), trophic level (FoodTroph, with DietTroph used as a fallback when FoodTroph was absent), feeding type (FeedingType field, referred to here as feeding guild), vertical position in the water column (DemersPelag), and a binary indicator of freshwater occurrence. Trait coverage across the 50 fish species was complete or near-complete (Figure 1; Table S1).

Four species lacked trophic level records; one received an imputed value equal to the genus-level median computed from all FishBase entries for that genus, while three could not be imputed as no congeneric FishBase entries were available and were therefore treated as missing. Nine species lacked a valid FishBase feeding guild category and could not be assigned one; these species were treated as having no guild match in pairwise feature construction. Non-fish nodes in the food webs (invertebrates, algae, detritus, and other basal resources) were assigned a trophic level of 1.0; all FishBase-specific fields were recorded as missing for these nodes. Prior to model fitting, missing body length values were filled with the within-fold training median (Petchey et al. 2008; Gravel et al. 2013). Fish species Lmax coverage was 100%, so this step applied only to non-fish basal nodes.

### 2.3 Cross-validation design

We evaluated all models using LOSO-CV. Networks were grouped by the study from which they originated, identified through the studyref metadata field in both databases. In each fold, all networks belonging to one study were withheld as the test set, and all networks from the remaining studies formed the training set. Each model was fitted exclusively on training-set data and then applied to all ordered node pairs in the held-out networks, with no information from the target system used at any stage of model fitting or feature construction. This procedure reflects the intended use case. It predicts the interaction structure of a food web in a system where no prior interaction data exist (Figure 2).

**Figure 2.**
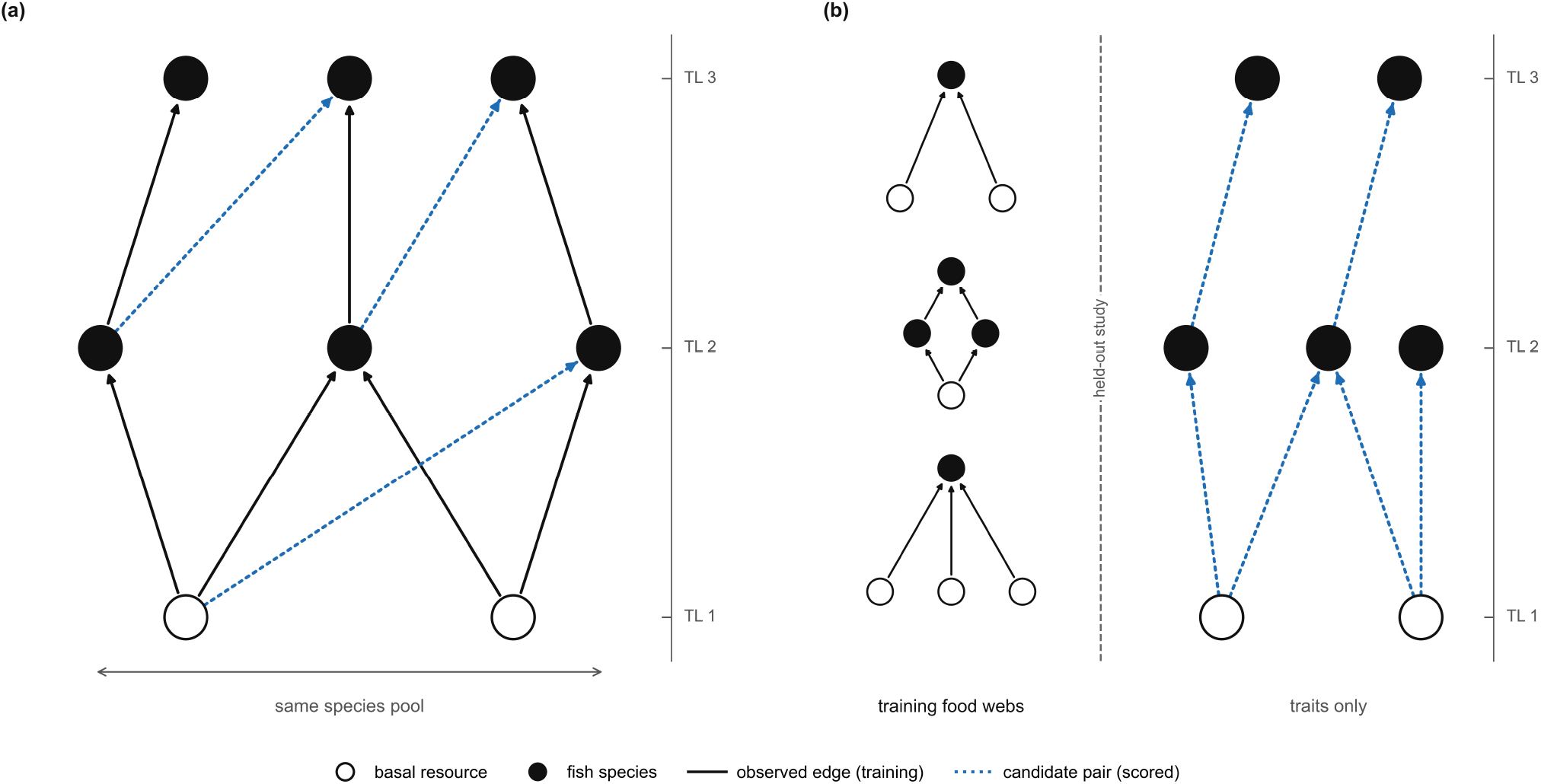
Evaluation designs compared in this study. (a) Within-network holdout: training edges (solid black) and held-out candidate pairs (blue dotted) are drawn from the same food web; the full species pool and all observed local interactions remain available to the model. (b) Leave-one-study-out transfer: models are trained on food webs from all other studies (solid black edges) and applied to every ordered consumer-resource pair in the held-out network using species traits only; no edges from the target network enter feature construction, model fitting, message passing, or prediction. Arrows indicate trophic direction (prey → predator). TL, trophic level.

In the GATEWAy database each network carries a unique study reference identifier, so LOSO-CV grouped by studyref produces one fold per network for all 34 GATEWAy entries. The three Mangal networks each carry a distinct study reference and constitute three additional folds. The full protocol therefore comprised 37 folds in total. The 27 Adirondack pelagic lake networks share a common geographic origin but differ in species composition and lake chemistry; each is treated as an independent fold, meaning the remaining 26 Adirondack lakes are available as training data when any one is held out.

We interpret results under two complementary evaluation perspectives. The first is within-source transfer: performance on the 34 GATEWAy folds, where the training and test networks share a common ecological region, trait measurement protocol, and database schema. The second is cross-source transfer: performance on the 3 Mangal networks, which were contributed independently by field researchers, span three continents, and have no overlap with the geographic or ecological composition of the GATEWAy training data. These networks were never used in training under any fold and represent the most demanding out-of-region test available in the current corpus.

### 2.4 Pairwise feature construction

For each network we enumerated all ordered node pairs (i, j) where i is the potential consumer and j is the potential resource, excluding self-loops. Pairs corresponding to observed edges were assigned a positive label (1); all other pairs received a negative label (0). The overall positive rate across all ordered node pairs in the 37 fish-rich networks was 14.4%, consistent with the high sparsity typical of empirical food webs. To address class imbalance during training, negative pairs were downsampled within each training network to five times the number of positive pairs (1:5 ratio; Pichler et al. 2020) before pooling across networks. At evaluation, all pairs in the held-out network were scored without any resampling so that performance metrics reflect the true, unbalanced class distribution. Three feature sets were constructed. Model A used three traits from GATEWAy: the natural logarithm of the consumer-to-resource body mass ratio (*x*_*a*_), a binary movement-type match indicator (*δ*^*mv*^), and a binary metabolic-type match indicator (*δ*^*met*^). Model B used eight FishBase-derived pairwise features. The primary size-asymmetry feature was the log Lmax ratio:

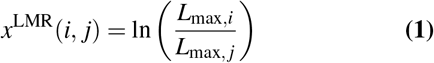

where *L*_max,*i*_ and *L*_max, *j*_ are the maximum body lengths (cm) of consumer i and resource j retrieved from FishBase. The trophic level difference was computed as:

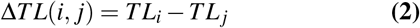

where *TL*_*i*_ and *TL*_*j*_ are the FoodTroph values for consumer i and resource j respectively. Additional features in Model B were binary indicators for feeding guild match (*δ* ^*gu*^), DemersPelag category match (*δ* ^*DO*^), and the absolute trophic levels and Lmax values of both nodes: *β* (*i, j*) = [*x*^*LMR*^, Δ*TL, TL*_*i*_, *TL*_*j*_ , *L*_max,*i*_, *L*_max, *j*_ , *δ*^*gu*^, *δ* ^*DO*^].

Model C extended Model B with a continuous habitat overlap score derived from the FishBase DemersPelag depth-stratum categories. Six categories were assigned ordinal depth ranks *d*_*k*_ ∈ 1, 2, …, 6 from shallowest to deepest (pelagic = 1, benthopelagic = 2, reef-associated = 3, demersal = 4, bathypelagic = 5, bathydemersal = 6). The overlap score was:

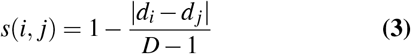

yielding s(i,j) = 1 when both nodes share the same stratum and s(i,j) = 0 when they occupy opposite ends of the depth gradient, where D = 6 is the number of depth categories.

### 2.5 Baseline and random forest models

Random forests (RF) were used for Models A, B, and C, implemented with scikit-learn (1.5.0; Pedregosa et al. 2011) in Python (3.11) using T = 300 trees, the square root of the number of features considered at each split, and a minimum of three samples per terminal leaf. A fixed random seed (42) was used throughout to ensure reproducibility. The predicted link probability for a consumer-resource pair (i, j) was the mean positive-class probability across all trees:

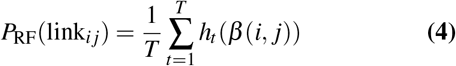

where *h*_*t*_(·) ∈ [0, 1] is the positive-class probability from tree t (Breiman 2001). Feature importance was quantified as mean decrease in impurity (MDI): the total decrease in node impurity (Gini index) weighted by the probability of reaching each node, averaged across all trees and normalised to sum to 1.0 (Breiman 2001). MDI is known to be biased toward high-cardinality continuous features and correlated predictors (Strobl et al. 2007); values should be interpreted with this limitation in mind.

Two baseline models were included to contextualise the performance of the learned models. Baseline 1 (size heuristic) predicted a link whenever the consumer’s maximum body length exceeded the resource’s (log Lmax ratio > 0), with no other information, establishing the predictive value of size rank alone. Baseline 2 (logistic regression) was fitted on the same eight FishBase pairwise features as Model B, providing a linear upper bound against which the non-linear RF could be assessed.

### 2.6 Graph attention network with inductive trait-only inference

Model D was a graph attention network (GAT; Velickovic et al. 2018) trained on food web structure across training networks. At inference on the held-out network, no observed edges are available; predictions are made from species trait vectors alone, without message passing.

#### 2.6.1 Architecture

The GAT comprised three components: (i) a trait encoder, (ii) two graph attention layers, and (iii) a bilinear link decoder. The trait encoder mapped per-node feature vectors to 32-dimensional initial embeddings via a single linear projection with exponential linear unit (ELU) activation. The 12-dimensional per-node input vector comprised normalised Lmax and trophic level, binary is-fish and is-basal flags, a 7-category one-hot encoding of DemersPelag vertical habitat stratum, and a normalised feeding guild index; these are per-node features distinct from the 8 pairwise features used in Models B and C.

The graph attention layers followed the original GAT formulation (Velickovic et al. 2018). The first layer used four attention heads with 8 dimensions each (output concatenated to 32 dimensions); the second used one head with 32 dimensions. Dropout (rate 0.3) and batch normalisation were applied after each layer. The attention coefficient for edge (i, j) in head k was computed as:

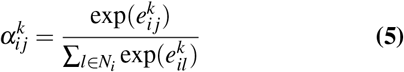

where 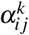 is the normalised attention weight for head k, 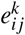 is the unnormalised attention score, and *N*_*i*_ is the neighbourhood of node i. The summation in the denominator runs over index l ∈ *N*_*i*_ to avoid conflict with the head index k (Velickovic et al. 2018). The node embedding update was:

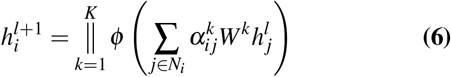

where K = 4 attention heads, *ϕ* is the ELU activation, ∥ denotes concatenation, and *W*^*k*^ is the head-specific weight matrix. A bilinear link decoder scored ordered node pairs from their learned embeddings:

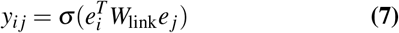

where *e*_*i*_ is the final node embedding, *W*_link_ is a learned 32× 32 weight matrix, and *σ* is the sigmoid function (Kipf and Welling 2016).

#### 2.6.2 Training procedure

In each LOSO fold, the GAT was trained on pooled training networks using the Adam optimiser (Kingma and Ba 2015; learning rate *η* = 0.001, weight decay *λ* = 0.0001) with a StepLR learning rate schedule (step size = 30, decay factor *γ* = 0.5). Training ran for a fixed 80 epochs; no validation-set early stopping was used, and the epoch count was determined from preliminary experiments monitoring training loss. Binary cross-entropy with logits loss was minimised, with the same 1:5 negative sampling per training network as for the RF. The GAT was implemented in PyTorch 2.1.0 (Paszke et al. 2019) with PyTorch Geometric 2.5.0 (Fey and Lenssen 2019). Full hyperparameter details are provided in Table S2.

#### 2.6.3 Inductive inference at test time

At test time, no edges from the held-out network were available. Because the graph attention layers require a known adjacency structure for message passing, they were bypassed entirely. The trait encoder mapped per-node feature vectors to embeddings directly, and the bilinear decoder scored all ordered pairs from those embeddings. No interaction data from the test network entered the model at any stage. This design tests whether structural regularities learned during training are captured in the trait encoder weights and yield useful predictions when no local interaction data exist.

### 2.7 Evaluation metrics and statistical comparison

Model performance was quantified using two threshold-independent metrics computed on all pairs in each held-out network: the area under the receiver operating characteristic curve (ROC-AUC) and the area under the precision-recall curve (PR-AUC). Both metrics were computed on the full, unbalanced test set using the trapezoidal rule. For a classifier operating at K thresholds with false positive rates FP*R*_*k*_ and true positive rates TP*R*_*k*_ (Fawcett 2006):

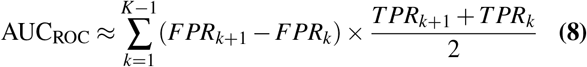

For the precision-recall curve, with precision *P*_*k*_ and recall *R*_*k*_ at threshold k (*R*_0_ = 0):

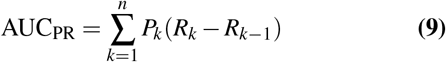

PR-AUC is given priority in our interpretation because it is more sensitive to performance on the minority positive class when class imbalance is substantial (Saito and Rehmsmeier 2015). A null classifier achieves PR-AUC approximately equal to the positive rate. This rate is 0.144 pooled across all ordered pairs in the corpus, but the median across the individual networks is 0.105. Because we summarise performance as the median across folds, we use 0.105 as the null for median PR-AUC comparisons. A null classifier achieves ROC-AUC = 0.50 regardless of class balance; PR-AUC is therefore the more stringent benchmark. Differences between models across LOSO folds were assessed using two-sided Wilcoxon signed-rank tests on paired per-network AUC values (Wilcoxon 1945), with significance threshold *α* = 0.05. Holm-Bonferroni correction was applied to control the family-wise error rate across the 12 pairwise comparisons; all significance decisions were unchanged after correction. Significance after Holm-Bonferroni correction is indicated in Table S3. Effect sizes were quantified as the median difference in AUC across folds. To locate the source of Model B’s full-network performance, we stratified each held-out GATEWAy test set into fish-associated pairs, in which the consumer or resource is a fish, and non-fish pairs, in which neither node is a fish, and recomputed ROC-AUC and PR-AUC within each class for Model A and Model B from the same full, unbalanced held-out predictions.

### 2.8 Software and reproducibility

All analyses were conducted in Python 3.11. Data processing used pandas 2.2.2 and NumPy 1.26.4. RF models were implemented with scikit-learn 1.5.0 (Pedregosa et al. 2011). The GAT was implemented in PyTorch 2.1.0 (Paszke et al. 2019) using PyTorch Geometric 2.5.0 (Fey and Lenssen 2019). Food web data were retrieved from GATEWAy via the GloBI GitHub repository and from Mangal using the rmangal package (3.0.0) in R 4.3.1. Species trait data were retrieved from FishBase using rfishbase 5.0 (Boettiger et al. 2012). Statistical tests (Wilcoxon signed-rank) were performed using scipy 1.13.0. Random seeds were fixed at 42 for Python and NumPy to ensure reproducibility.

## Results

### 3.1 Corpus characteristics

The 34 GATEWAy networks included 27 Adirondack pelagic lakes, 6 English stream systems, and 1 New Zealand stream (Dempsters Stream). The 3 Mangal networks comprise Lake Crescent (Washington, USA), Gatun Lake (Panama) and Potreirinho Creek (Brazil), representing the geographically distinct out-of-region test cases. Full metadata for all 37 networks are provided in Table S4.

### 3.2 Model A: locally measured body mass ratios achieve near-perfect prediction

Model A, trained on three GATE-WAy traits (log body mass ratio, movement type match, and metabolic type match) and evaluated on GATEWAy fish-rich networks only, demonstrated that trophic links are highly predictable when locally measured body mass data are available. Across the 34 networks (n = 34 LOSO folds), the median ROC-AUC was 0.973 (interquartile range 0.941 to 0.978) and the median PR-AUC was 0.829 (IQR: 0.754 to 0.871; Table 1). Performance was consistently high: 24 of the 34 networks achieved a ROC-AUC at or above 0.95, and only 7 networks fell below 0.90 (Figure 3). The matched panels show how link recovery changes when the same held-out network is predicted from locally measured GATEWAy body-mass features versus FishBase-derived traits, for Hoel Lake (Figure 4a,b), Long Lake (Figure 4c,d), and Mill Stream (Figure 4e,f). Log body mass ratio accounted for more than 95% of mean feature importance in Model A, confirming that predator-prey size asymmetry is the dominant signal in these networks. Because GATEWAy body mass data are unavailable for the Mangal networks, Model A serves as a within-database reference baseline and is not included in cross-source comparisons.

**Table 1.** Leave-one-study-out cross-validation performance for all four models.

| Model | Feature set | n | ROC-AUC | PR-AUC |
| --- | --- | --- | --- | --- |
| Full corpus (n = 37) |  |  |  |  |
| Model B | FishBase: 8 pairwise features (Lmax, trophic level, feeding guild, DemersPelag) | 37 | 0.607 | 0.207 |
| Model C | Model B + habitat overlap score (9 features) | 37 | 0.606 | 0.206 |
| Model D | FishBase 8 traits; inductive trait-only inference (GAT) | 37 | 0.554 | 0.144 |
| GATEWAY networks (n = 34) |  |  |  |  |
| Model A | Local body mass: log mass ratio, movement match, metabolic match | 34 | 0.973 | 0.829 |
| Model B | FishBase 8 pairwise features | 34 | 0.609 | 0.215 |
| Model C | Model B + habitat overlap (9 features) | 34 | 0.606 | 0.213 |
| Model D | FishBase 8 traits, inductive trait-only inference (GAT) | 34 | 0.553 | 0.144 |

Note. ROC-AUC and PR-AUC are medians across LOSO folds. Null classifier: ROC-AUC = 0.500; PR-AUC equals the positive rate (median per-network 0.105; 0.144 pooled across all pairs). Full pairwise Wilcoxon comparisons in Table S3.

**Figure 3.**
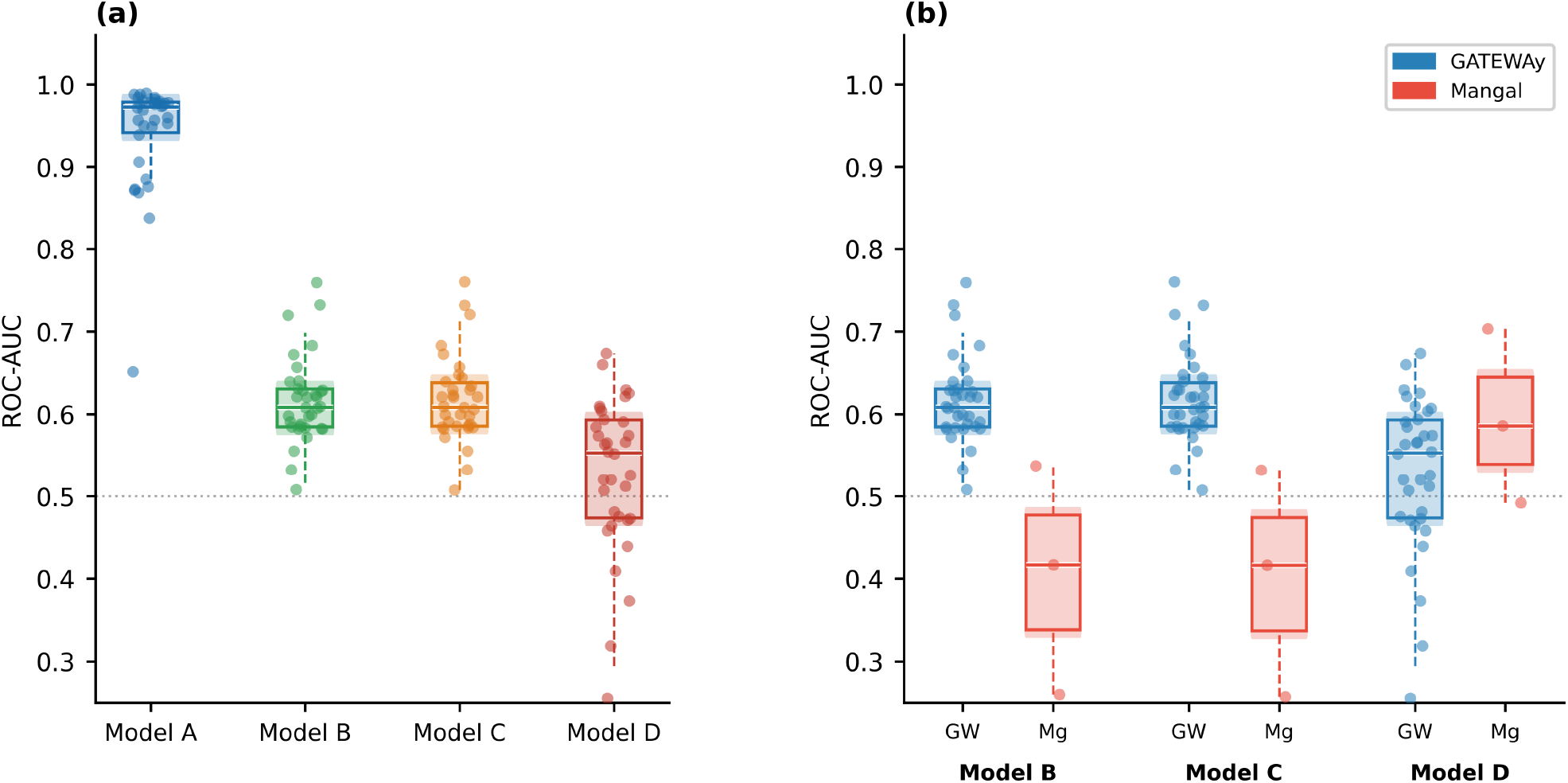
Leave-one-study-out cross-validation ROC-AUC for all four models. (a) Distribution across the 34 GATEWAy networks for Models A, B, C, and D. (b) Distribution by source database for Models B, C, and D across all 37 networks. GW = GATEWAy; Mg = Mangal. Boxes show interquartile range; white bars show medians; points are individual networks (jittered). Dotted horizontal line marks chance level (ROC-AUC = 0.50). PR-AUC results are shown in Figure S1.

**Figure 4.**
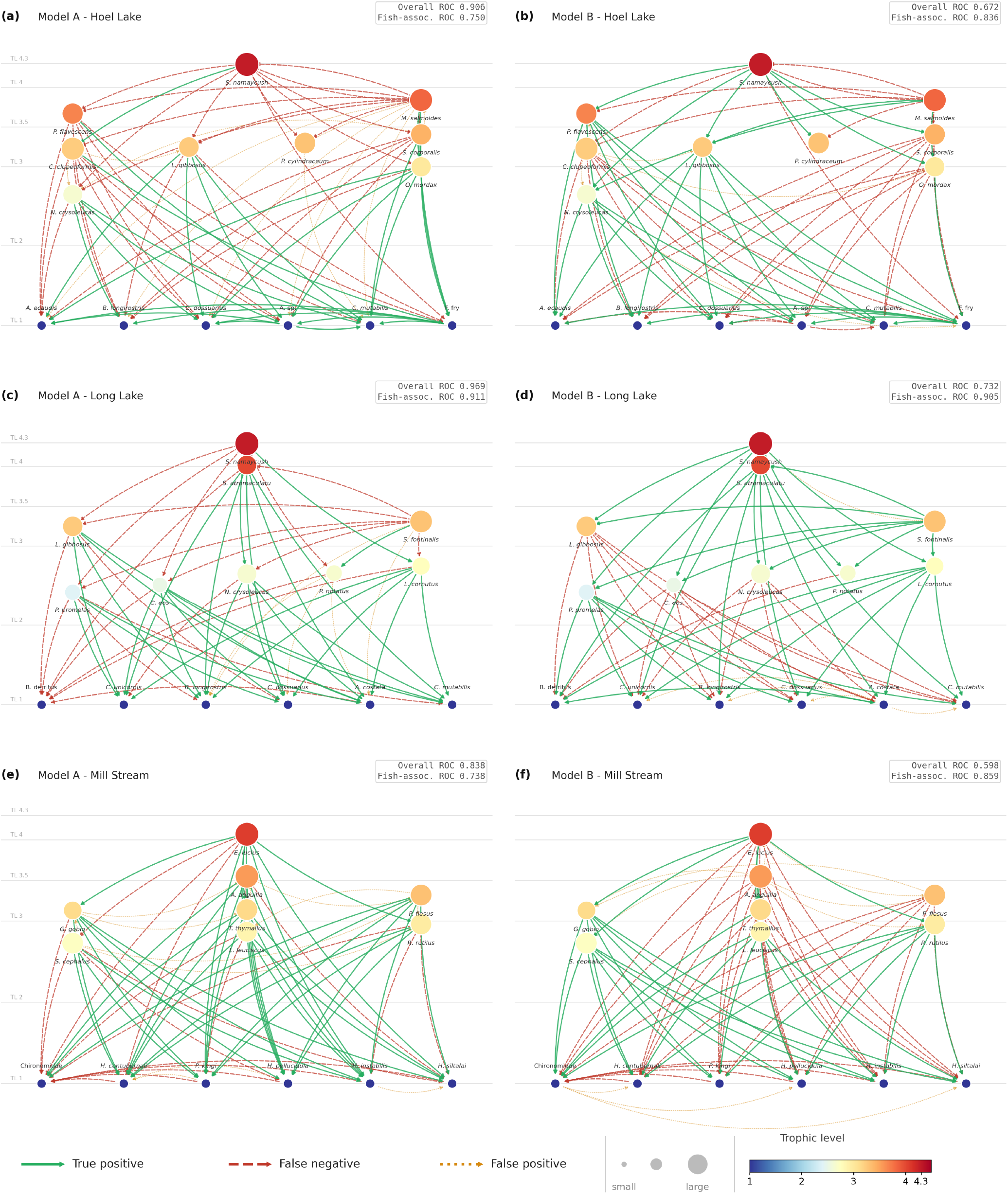
Matched Model A and Model B link predictions in three held-out networks. Each row is one GATEWAy network held out under leave-one-study-out cross-validation, with Model A (locally measured GATEWAy body mass) on the left and Model B (FishBase-derived traits) on the right, drawn on the same nodes. Nodes are positioned and coloured by trophic level and sized by FishBase maximum body length on a shared scale across panels. Green solid arrows are observed links recovered by the model (true positives), red dashed arrows are observed links missed (false negatives), and orange dotted arrows are unobserved pairs predicted as links (false positives). Each panel label gives ROC-AUC over all ordered pairs in the full held-out network (Overall) and over fish-associated pairs only (Fish-assoc.). Panels show (a, b) Hoel Lake, (c, d) Long Lake, and (e, f) Mill Stream, the three most fish-rich networks in the corpus. Across the three networks the overall ROC-AUC falls from Model A to Model B; the fish-associated ROC-AUC does not.

### 3.3 Models B and C: FishBase trait-based cross-system prediction

Model B (eight FishBase traits) achieved a median ROC-AUC of 0.607 (IQR: 0.583 to 0.629) and a median PR-AUC of 0.207 (IQR: 0.173 to 0.249) across all 37 networks. Model C (Model B plus habitat overlap score) achieved a median ROC-AUC of 0.606 and a median PR-AUC of 0.206. Wilcoxon signed-rank tests returned no significant difference between Models B and C for either metric (ROC-AUC: W = 89.0, p = 0.356; PR-AUC: W = 68.0, p = 0.167, Holm-adjusted 0.334; Table S3). The habitat overlap feature contributed a mean importance weight of only 0.006, the lowest of any feature in either model (Table 2), indicating that FishBase depth-stratum categories carry no usable signal for local trophic prediction.

**Table 2.** Mean feature importance for Models B and C, averaged across leave-one-study-out folds.

| Feature | Imp. B | Imp. C | Description |
| --- | --- | --- | --- |
| Trophic level difference | 0.240 | 0.245 | Consumer minus resource trophic level |
| Consumer Lmax (cm) | 0.230 | 0.236 | Max body length of consumer (FishBase) |
| Resource Lmax (cm) | 0.201 | 0.207 | Max body length of resource (FishBase) |
| Consumer trophic level | 0.166 | 0.160 | Consumer trophic level (FoodTroph) |
| Resource trophic level | 0.075 | 0.074 | Resource trophic level (FoodTroph) |
| Feeding guild match | 0.037 | 0.025 | Consumer and resource share feeding guild |
| DemersPelag match | 0.035 | 0.032 | Consumer and resource share depth stratum |
| Log Lmax ratio | 0.016 | 0.015 | $\ln(\text{consumer Lmax} / \text{resource Lmax})$ |
| Habitat overlap score | n/a | 0.006 | Depth-stratum overlap score (0-1; Model C only) |

*Note.* Imp. = mean decrease in impurity (MDI), normalised to sum to 1.0, averaged across 300 trees and all LOSO folds. n/a = feature not included in this model. Ranked by Model B importance.

Performance differed markedly between databases. On GATE-WAy networks, Model B achieved median ROC-AUC 0.609 and PR-AUC 0.215, substantially below Model A despite identical network coverage. Splitting each held-out network by pair type located most of this gap (Figure 5; Table S5). For fish-associated pairs, in which the consumer or resource is a fish, the two models were close (Model A median ROC-AUC 0.912, Model B 0.887). The difference between them fell almost entirely on pairs between non-fish nodes, where Model A stayed high (0.979) but Model B was at chance (0.500), since FishBase records no traits for these taxa. Non-fish pairs form the majority in each network, so they set the level of the full-network median. Performance on the three Mangal networks was lower (median ROC-AUC 0.417, PR-AUC 0.127). With only three out-of-region networks, these results are exploratory and do not support general conclusions (Table 1; Figure 7). The pair-class split did not recover performance there either. Fish-associated ROC-AUC did not reach the within-GATEWAy level, ranging from below chance to marginally above it across the three networks.

**Figure 5.**
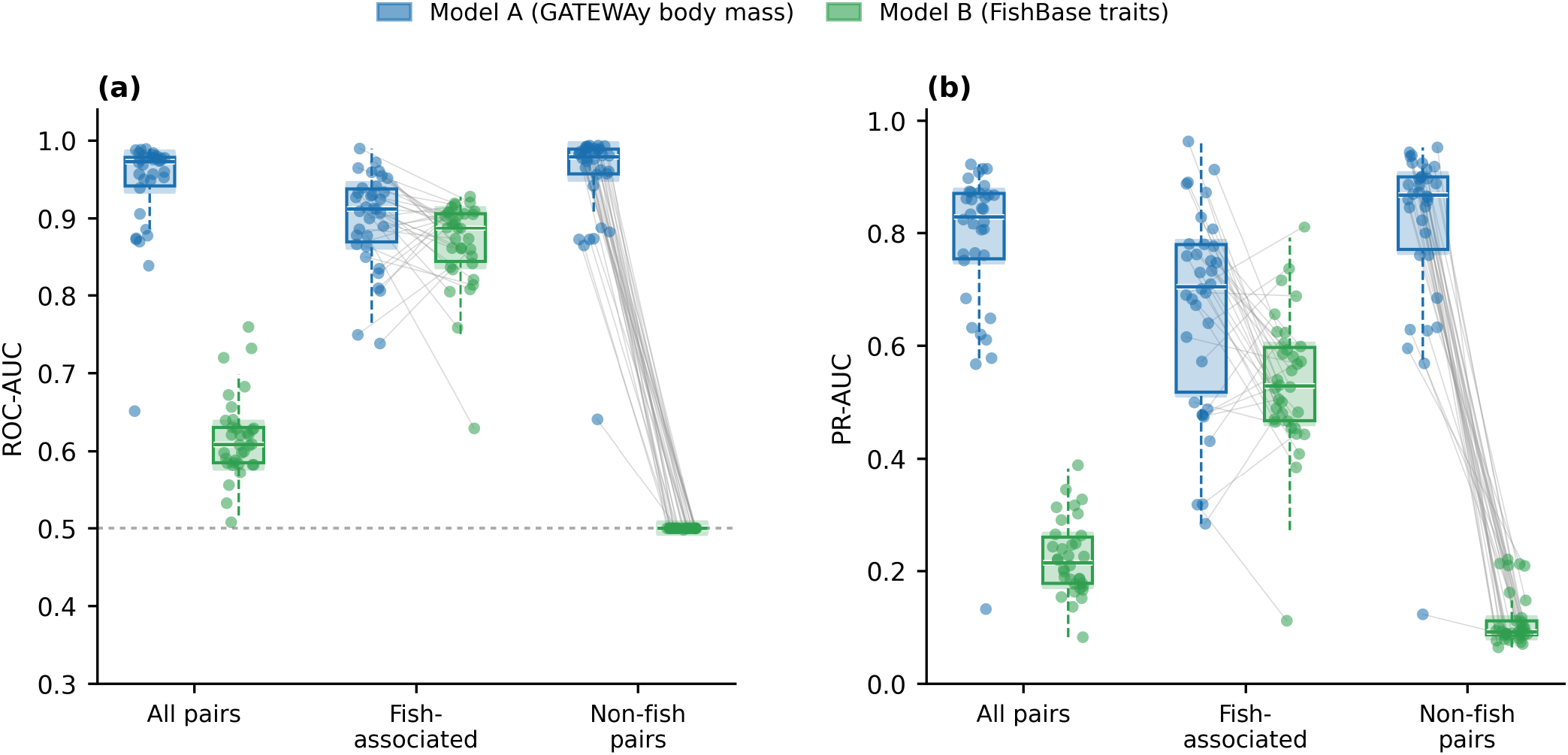
Model A and Model B performance by pair class across the 34 GATEWAy networks. (a) ROC-AUC and (b) PR-AUC. Points are individual networks; boxes show the interquartile range and white bars the medians. Fish-associated pairs are ordered pairs in which the consumer or resource is a fish; non-fish pairs are pairs in which neither node is a fish. Model A used locally measured GATEWAy body mass, Model B used FishBase-derived traits. The dashed line marks chance (ROC-AUC = 0.50).

Baseline 2, the logistic regression trained on the same eight FishBase features, achieved median ROC-AUC of 0.607 across all 37 networks, matching Model B exactly. However, its median PR-AUC was 0.193 compared to 0.207 for the RF (Wilcoxon signed-rank W = 98, p < 0.001, Holm-adjusted p < 0.001). The RF outperformed logistic regression in 29 of 37 networks on PR-AUC, indicating that it captures non-linear feature interactions that improve recovery of positive links under class imbalance. Baseline 1, the size heuristic (consumer Lmax > resource Lmax) was near-random, achieving median ROC-AUC of 0.502 and median PR-AUC of 0.109 across all 37 networks, at the chance level set by the median per-network positive rate (0.105). Size rank alone therefore provides negligible predictive power, and the gains achieved by Models B and C reflect the information contributed by trophic level, feeding guild, and their pairwise combinations rather than any simple size-ordering rule.

### 3.4 Model D: graph-structured training does not improve zero-edge cold-start prediction overall

Model D (GAT with FishBase traits) achieved a median ROC-AUC of 0.554 and PR-AUC of 0.144 overall, significantly below Model B (ROC-AUC: W = 80.0, p < 0.001; Table S3). Model D scored higher than Model B on the three Mangal networks (median ROC-AUC 0.586 versus 0.417). This difference comes from only three out-of-region networks, one of which, Lake Crescent, has seven directed edges, and we did not isolate its cause with an ablation experiment (Table 1; Figure 3b).

### 3.5 Relative importance of trait features

In both Models B and C, trophic level difference between consumer and resource was the most important feature, accounting for 24.0% of mean feature importance in Model B (Table 2). The two absolute body size features (consumer Lmax and resource Lmax) together accounted for approximately 43% of importance in both models. In contrast, the log Lmax ratio ranked last among continuous features with a mean importance of only 1.6% (Figure 6a). Permutation importance analysis, computed on held-out test networks per fold, placed resource trophic level first in Model B (mean decrease in ROC-AUC = 0.010) and trophic level difference first in Model C (0.009; Table S6). Consumer and resource Lmax ranked lower under permutation than under MDI, consistent with known MDI inflation of high-cardinality continuous features (Strobl et al. 2007). Per-network detail is provided in Figures 7, S2, S3, and S4.

**Figure 6.**
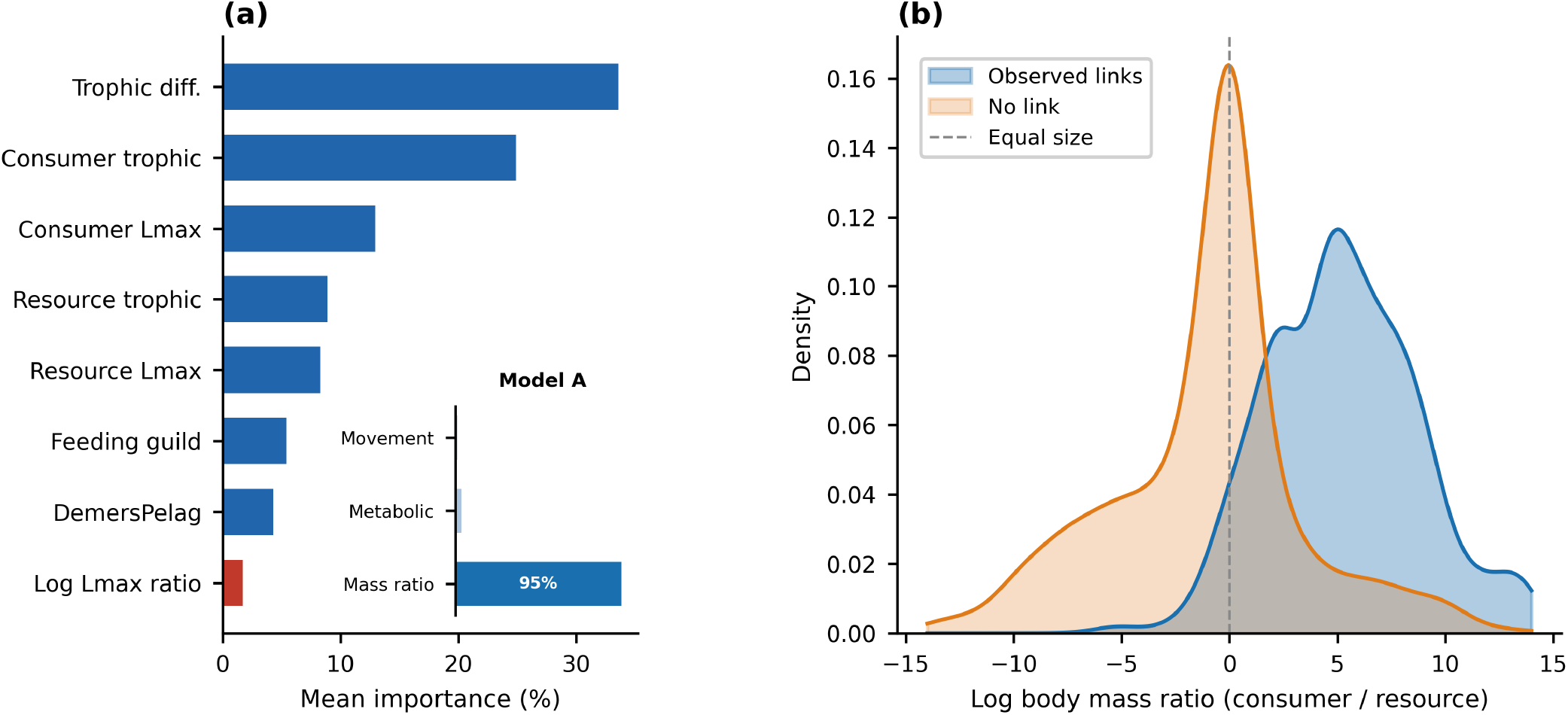
Feature contributions and trait-based separability. (a) Mean feature importance (%) for Model B (main bars) and Model A (inset). Log Lmax ratio (red) is the Model B analogue of the body mass ratio that dominates Model A importance (>95%). (b) Kernel density distributions of log body mass ratio (natural log of consumer mass divided by resource mass) for observed trophic links (blue) and absent links (orange) in the GATEWAy dataset. Dashed vertical line marks equal consumer and resource body mass (ratio = 1). The x-axis covers the 5th-95th percentile of the data.

**Figure 7.**
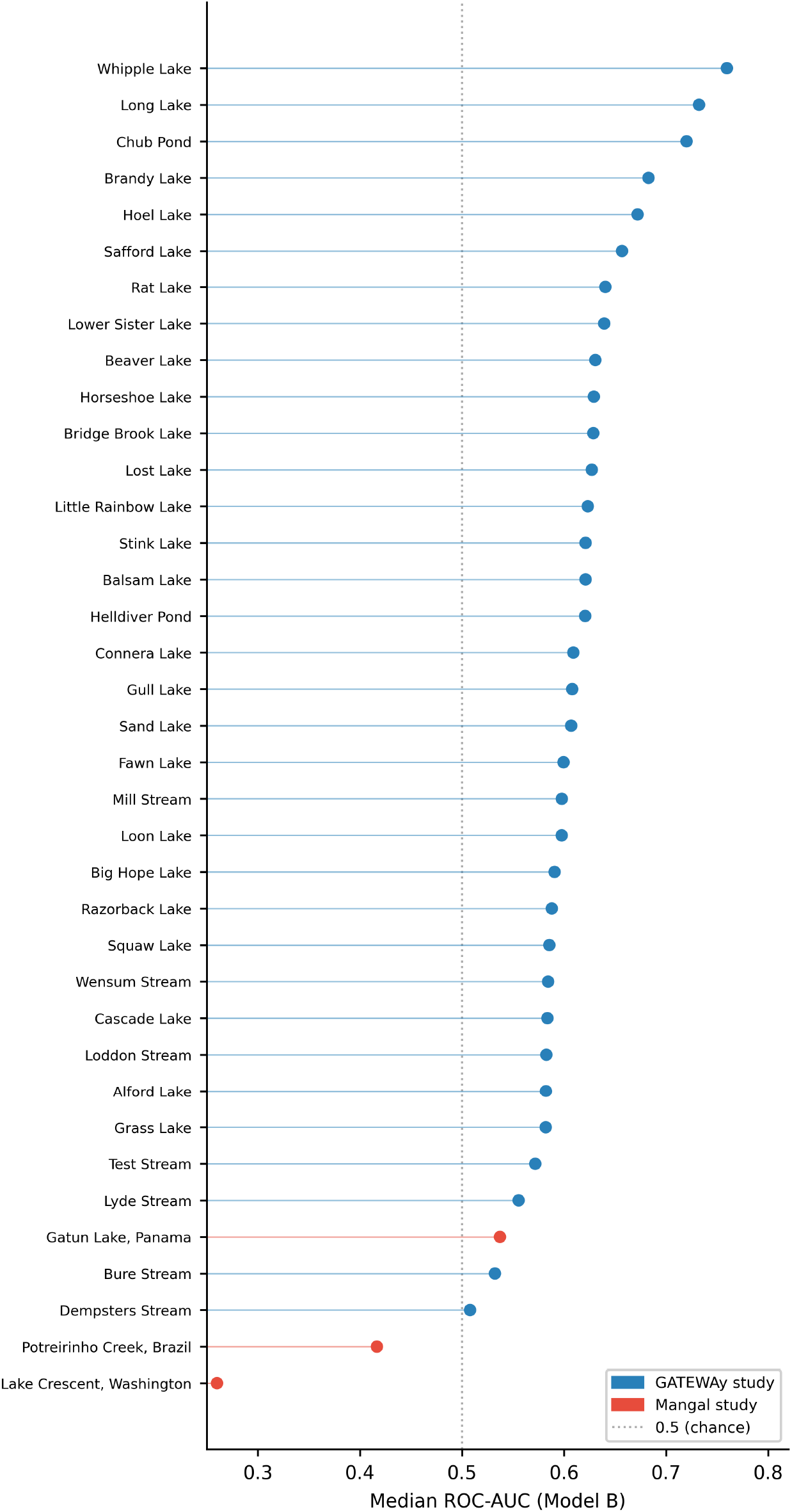
Per-network ROC-AUC for Model B across all 37 networks. Each point represents one network. Dotted vertical line marks chance level (ROC-AUC = 0.50). Blue = GATEWAy networks; red = Mangal networks. Per-network PR-AUC results are shown in Figure S2.

## Discussion

### 4.1 Coverage and precision in the cross-system gap

Our results confirm that body mass asymmetry is the dominant organising trait in these networks (Brose et al. 2006; Woodward et al. 2005; Petchey et al. 2008; Gravel et al. 2013), but add a critical qualification. The predictive value of body size depends entirely on which size metric is available. When locally measured body mass ratios were used, the RF achieved median ROC-AUC of 0.973 across 34 networks, with body mass ratio accounting for more than 95% of mean feature importance (Figure 6a). The same algorithm, applied to the same networks with FishBase maximum body length substituted for local body mass, reached only 0.609 across the same 34 networks. The pair-class comparison shows where this gap comes from (Figure 5; Table S5). For fish-associated pairs, Model B nearly matched Model A (0.887 against 0.912); the coarse global size measure lost little signal. For pairs between non-fish nodes, Model A stayed accurate (0.979) while Model B did not, because FishBase records no traits for these taxa. Non-fish pairs make up most of each web and set the whole-network score. FishBase traits predict the fish interactions this study targets. Only three networks lie outside a single temperate region, so out-of-region performance is not yet established.

The reasons for this gap bear directly on how researchers should collect and use trait data. FishBase Lmax records the largest individual ever documented for a species across its full geographic range (Froese and Pauly 2026). It is not an estimate of typical adult size at a particular location, nor does it capture the size structure of the population present in a given lake or stream at the time of sampling. A juvenile and an adult of the same species carry identical Lmax values but occupy different trophic positions (Werner and Gilliam 1984). A population of large-bodied trout in an oligotrophic lake and a stunted population of the same species in an acidified lake receive the same Lmax record. The consumer-to-resource body mass ratio that drives Model A’s near-perfect predictions cannot be reconstructed from these range-wide extremes. On these fish-associated pairs, the signal came from trophic level and feeding guild, not from the size ratio.

That the log Lmax ratio ranked last among continuous features in Models B and C (mean importance 1.6%; Table 2) confirms that this is a signal loss problem rather than an algorithmic one (Table 2; Table S6; Figure 6a). Permutation importance placed trophic level features above the size ratio (Table S6), suggesting that when the body size signal is degraded by a coarse global proxy, trophic level becomes the most informative available predictor because it is more consistently defined across species and less sensitive to local population structure. This shift in feature importance reveals what the model falls back on when its primary predictor fails. Consumer trophic level showed slightly negative permutation importance in Model B (Table S6), meaning that permuting it marginally improved hold-out performance on average. Consumer trophic level and trophic level difference are strongly collinear. Permuting one changes the effective contribution of the other, so the negative importance is an artefact of correlated predictors rather than a true lack of information (Strobl et al. 2007). A linear classifier on the same FishBase features matched the RF exactly in ROC-AUC (both 0.607), confirming that the bottleneck is the feature set, not model complexity. Better body size data would benefit any classifier.

### 4.2 The habitat information problem

The habitat overlap score derived from FishBase DemersPelag depth-stratum categories added no predictive value beyond the eight features already in Model B (Tables 2 and S3). This null result reveals a structural limitation of global trait databases that extends beyond body size.

The DemersPelag field in FishBase describes where a species has been recorded across its entire geographic range (Froese and Pauly 2026). It captures a biogeographic tendency, not local foraging behaviour. In a shallow, well-mixed lake, species classified as pelagic and demersal may occupy the same water column throughout the day. The overlap score therefore measures geographic-scale vertical zonation rather than local spatial co-occurrence. This mismatch parallels the body size problem but arises through a different mechanism. Lmax fails because it ignores local population structure, while DemersPelag fails because it conflates biogeographic range with local behaviour.

This does not mean that habitat information is inherently uninformative for trophic prediction. Wootton et al. (2022) showed that microhabitat overlap improved predictions in a dynamic food-web model tested under controlled mesocosm conditions, and our own Model A results show that movement-type and metabolic-type matches carry signal when derived from local observations. The specific habitat categorisation available in FishBase is simply too coarse and too decoupled from local foraging behaviour to contribute usable signal at the community scale. Whether finer-grained environmental data from remote sensing or locally measured habitat use traits could add the predictive value that global database categories lack deserves testing.

### 4.3 Limits of graph-structured training in zero-edge inference

The GAT’s overall underperformance relative to the RF (ROC-AUC 0.554 versus 0.607; Table 1) reflects the constraints of zero-edge inductive inference. With no edges available in the held-out network, message passing was bypassed at test time, capping what graph-structured training could add to link prediction. The higher Model D scores on the three Mangal networks are exploratory because they rest on only three out-of-region tests and no ablation isolates a structural cause. Graph models may become more useful when partial target-network structure is available or when comparable trait coverage extends to more taxa.

### 4.4 Geographic bias and the limits of generalization

The most consequential limitation of this study is the composition of the training corpus. Twenty-seven of the 34 GATEWAy networks originate from a single regional survey of Adirondack lakes. These folds are therefore not fully independent replicates. The three Mangal networks are the only out-of-region evaluation cases, and they represent just three data points. Africa, southern Asia, tropical South America, and the Southern Hemisphere are entirely absent. The 50 fish species in this corpus represent a small fraction of the roughly 15,000 described freshwater species (Levêque et al. 2008). Poisot et al. (2021) documented the geographic bias in food web data toward temperate North American and European systems, noting that the regions most underrepresented in the literature are also those where the most divergent ecological contexts occur. This geographic concentration limits the ecological range represented in training, while the three Mangal folds are too few to support a site-specific mechanism.

Caron et al. (2024) reported a similar pattern in terrestrial vertebrate food webs, finding that trait-based model performance declined systematically with environmental and phylogenetic distance between training and test systems. Their ROC-AUC values remained above 0.75 even for the most distant transfers, likely because vertebrate body size is more reliably estimated from museum specimens than fish maximum length is from occurrence records. This comparison highlights that the severity of the trait-proxy problem depends on the taxon and the trait database. For fish, Lmax is particularly coarse because it records a range-wide extreme rather than a typical adult size, and because fish body size varies substantially with local environmental conditions such as temperature and productivity (Belk and Houston 2002).

The composition of the training corpus is itself an analytical decision. For a target ecosystem of known type, filtering or weighting training networks toward ecologically similar systems, even at the cost of fewer training networks overall, is likely to outperform simply pooling all available data regardless of ecological similarity. Hirn et al. (2024) demonstrated a related principle for transfer learning of species co-occurrence patterns between plant communities, finding that matching species composition between source and target communities improved transfer accuracy, particularly for small target datasets. This similarity-weighted training approach could be tested with the corpus that already exists, though its full value would be realised only with a substantially larger and more ecologically diverse collection of food webs spanning the full range of freshwater ecosystem types that practitioners need to predict.

The models in this study predict binary interaction presence or absence, not interaction strength. Empirical food webs vary substantially in the frequency and energetic importance of their links, and a model that correctly predicts which links exist but misses their relative importance may still provide incomplete guidance for management applications. Additional traits such as diet breadth, gape morphology, or swimming speed might improve prediction but are not yet available at the global scale for most freshwater fish (Wootton et al. 2022; Petchey et al. 2008; Caron et al. 2022). Incorporating phylogenetic distance from resources such as the Fish Tree of Life (Rabosky et al. 2018) as a complement to functional trait similarity could add discriminating power in systems where convergent morphology between ecologically distinct lineages weakens trait-based prediction. Strydom et al. (2023) showed that phylogenetic transfer of network representations can partially compensate for missing trait information, and combining their approach with the trait-based models tested here deserves exploration.

## Conclusions

Our results show that trophic links in freshwater fish food webs are highly predictable from traits when locally measured body mass data are available. Substituting FishBase maximum body length as a global proxy reduces median ROC-AUC from 0.973 to 0.609 across the same 34 GATEWAy networks under an otherwise identical evaluation design. This drop is confined to pairs between non-fish taxa, which FishBase does not describe. For pairs involving a fish, FishBase traits matched local body mass (0.887 against 0.912). A linear classifier fitted on the same FishBase features matched the RF’s ROC-AUC exactly, with only a modest gap in PR-AUC. Habitat overlap scores derived from FishBase depth-stratum categories added no predictive value, confirming that global-range categorisations do not capture local foraging behaviour. Within the ecological range of the training data, the practical implication is clear. Recording mean adult body mass per species per site during food web sampling is the most effective change to field protocols for improving cross-system prediction accuracy. The pair-class analysis clarifies where the loss comes from. FishBase traits retain substantial signal for fish-associated interactions; non-fish pairs are effectively uninformative because comparable trait data are absent. FishBase traits already predict fish-associated links in unstudied systems nearly as well as locally measured body mass. Wider prediction depends on assembling more fish food webs across more regions.

## DATA AVAILABILITY

Raw food web data were obtained from the GATEWAy database (Global Archive of Trait, Evolution, and Energy Use; Brose et al. 2019; https://github.com/globalbioticinteractions/brose-gateway) and Mangal (https://mangal.io; Poisot et al. 2016). Species trait data were retrieved from Fish-Base (https://www.fishbase.org; Froese and Pauly 2026) using rfishbase 5.0 in R 4.3.1.

## CODE AVAILABILITY

Analysis code used to construct features, fit models, evaluate performance, and generate figures is available at https://github.com/imranbinyounos/freshwater-fish-foodweb-link-prediction.

## AUTHOR CONTRIBUTIONS

Imran Bin Younos: Conceptualization, data curation, methodology, software, formal analysis, visualization, writing – original draft.

## ACKNOWLEDGEMENTS

I thank the curators of the GATEWAy database (Brose et al. 2019), the Mangal network repository (Poisot et al. 2016), and FishBase (Froese and Pauly 2026) for making their data publicly available.

