## Supplementary Information for "Predicting trophic links across freshwater fish food webs from species traits"

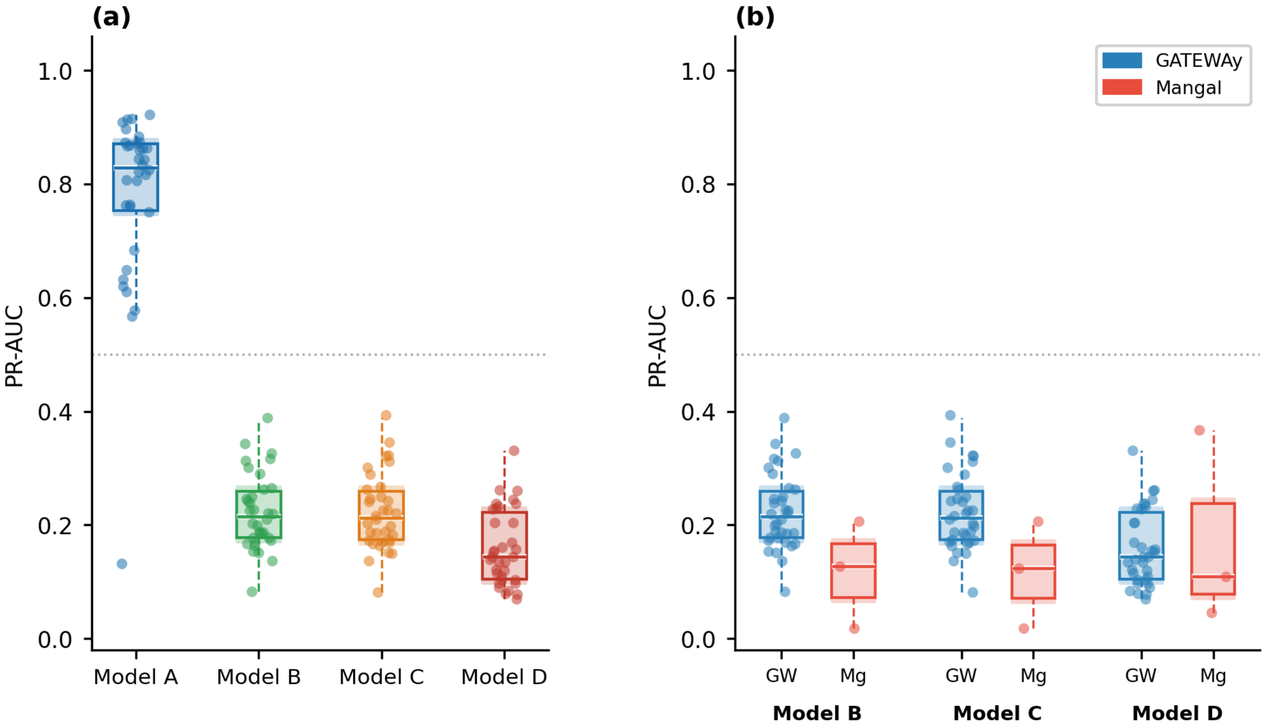


**Supplementary Figure S1.** Leave-one-study-out cross-validation PR-AUC for all four models. (a) Distribution across the 34 GATEWAy networks. (b) Distribution by source database for Models B, C, and D across all 37 networks. PR-AUC is weighted more heavily in interpretation than ROC-AUC because it is more sensitive to classifier performance on the minority positive class given the 14.4% positive rate of this corpus. A null classifier achieves PR-AUC approximately equal to the positive rate (0.144).

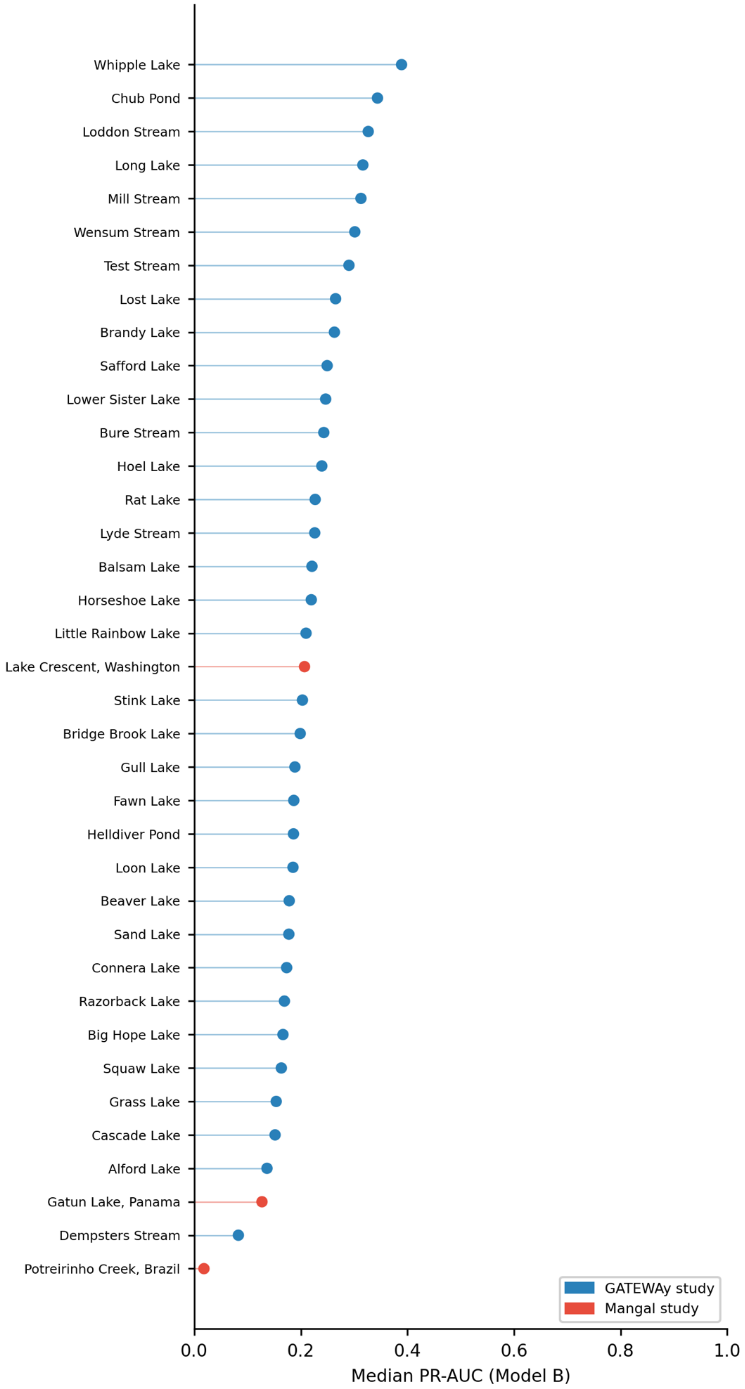


**Supplementary Figure S2.** Per-network PR-AUC for Model B across all 37 networks. Each point represents one network. Blue = GATEWAy networks; red = Mangal networks.

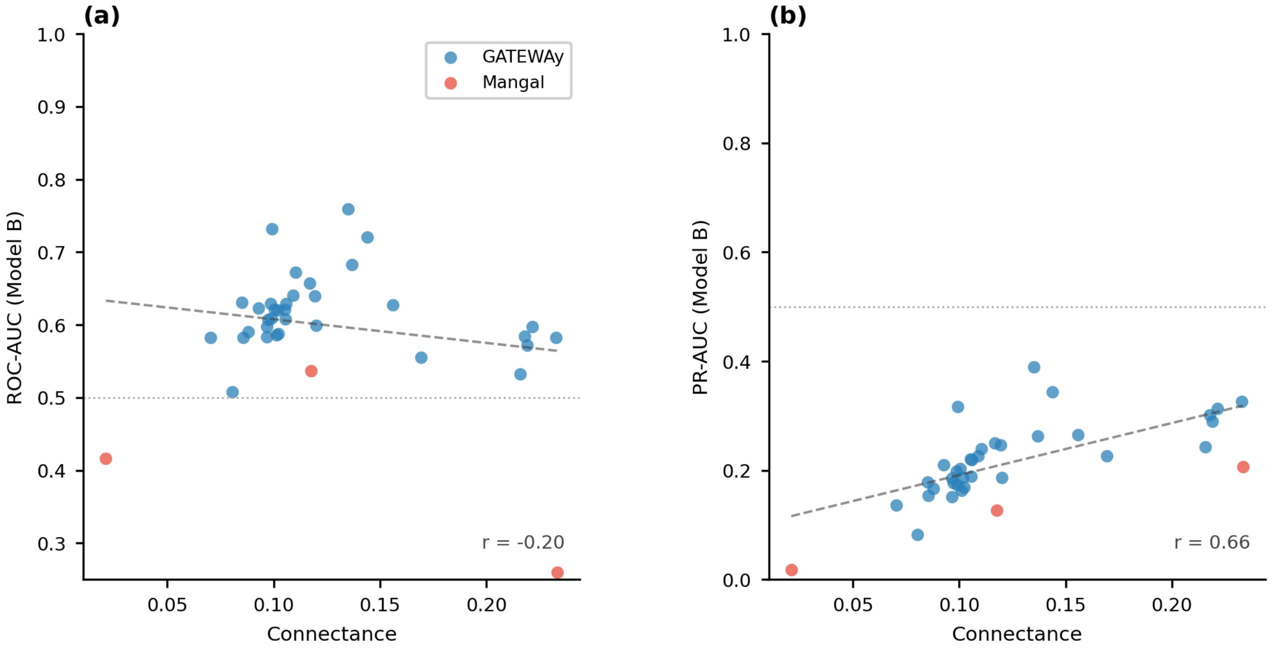


**Supplementary Figure S3.** Relationship between network connectance and Model B prediction performance. Each point represents one network; connectance is computed as observed edges divided by n(n-1) directed pairs. (a) ROC-AUC; (b) PR-AUC. Dashed lines show ordinary least squares fits across all 37 networks. Pearson correlation coefficients are shown in each panel. Dotted horizontal lines mark chance level (ROC-AUC = 0.50; PR-AUC = 0.144).

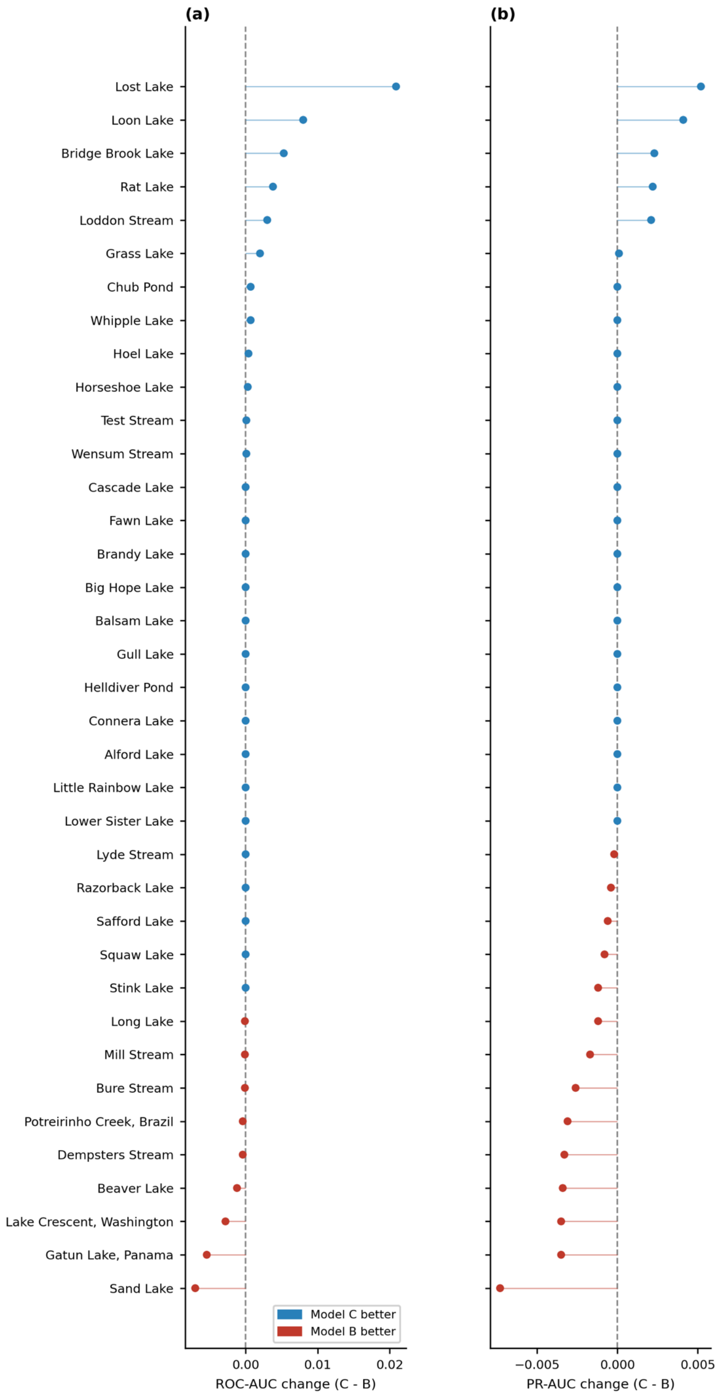


**Supplementary Figure S4.** Per-study mean difference in ROC-AUC (a) and PR-AUC (b) between Model C and Model B (positive values indicate Model C outperforms Model B). The 27 Adirondack pelagic lake networks and 6 English stream networks and 1 New Zealand stream network (Dempsters Stream) are each shown as their per-study mean delta. The 3 Mangal networks (Lake Crescent, Gatun Lake, Potreirinho Creek) are shown individually. Blue = Model C better; red = Model B better. Dashed vertical line marks zero difference.

**Table S1.** FishBase trait records for the 50 fish species included in the corpus.

| Species | Lmax (cm) | Trophic level | Feeding guild | DemersPelag | Family | n netw. |
|---|---|---|---|---|---|---|
| *Ambloplites rupestris* | 43.0 | 3.43 | hunting macrofauna | benthopelagic | Centrarchidae | 1 |
| *Anguilla anguilla* | 121.5 | 3.55 | hunting macrofauna | demersal | Anguillidae | 3 |
| *Anguilla dieffenbachii* | 185.0 | 4.47 | hunting macrofauna | demersal | Anguillidae | 1 |
| *Astyanax ruberrimus* | 2.6 | 3.18* | – | benthopelagic | Acestrorhamphidae | 1 |
| *Astyanax scabripinnis* | 15.6 | 3.18 | –* | benthopelagic | Acestrorhamphidae | 1 |
| *Barbatula barbatula* | 21.0 | 3.28 | hunting macrofauna | demersal | Nemacheilidae | 4 |
| *Catostomus catostomus* | 64.0 | 2.54 | grazing aquatic plants | demersal | Catostomidae | 2 |
| *Characidium schubarti* | 51.3 | –* | –* | benthopelagic | Crenuchidae | 1 |
| *Chrosomus eos* | 8.0 | 2.51 | variable | demersal | Leuciscidae | 12 |
| *Compsura gorgonae* | 2.9 | –* | –* | benthopelagic | Characidae | 1 |
| *Coregonus clupeaformis* | 100.0 | 3.23 | hunting macrofauna | benthopelagic | Salmonidae | 2 |
| *Cottus gobio* | 18.0 | 3.23 | hunting macrofauna | demersal | Cottidae | 4 |
| *Esox lucius* | 137.0 | 4.07 | hunting macrofauna | pelagic | Esocidae | 1 |
| *Fundulus diaphanus* | 13.0 | 3.30 | variable | benthopelagic | Fundulidae | 4 |
| *Galaxias depressiceps* | 8.2 | 3.21 | hunting macrofauna | benthopelagic | Galaxiidae | 1 |
| *Gasterosteus aculeatus* | 11.0 | 3.31 | hunting macrofauna | benthopelagic | Gasterosteidae | 4 |
| *Geophagus brasiliensis* | 28.0 | 2.57 | –* | benthopelagic | Cichlidae | 1 |
| *Gobio gobio* | 21.0 | 3.13 | hunting macrofauna | benthopelagic | Gobionidae | 2 |
| *Gobiomorphus breviceps* | 8.2 | 3.24 | –* | demersal | Eleotridae | 1 |
| *Gobiomorphus dormitor* | 90.0 | 3.63 | hunting macrofauna | demersal | Eleotridae | 1 |
| *Hoplias malabaricus* | 65.0 | 4.50 | hunting macrofauna | benthopelagic | Erythrinidae | 1 |
| *Imparfinis mirini* | 8.5 | 3.24 | –* | demersal | Heptapteridae | 1 |
| *Lepomis gibbosus* | 40.0 | 3.25 | hunting macrofauna | benthopelagic | Centrarchidae | 24 |
| *Leuciscus leuciscus* | 40.0 | 2.87 | hunting macrofauna | benthopelagic | Leuciscidae | 1 |
| *Luxilus cornutus* | 18.0 | 2.75 | variable | demersal | Leuciscidae | 6 |
| *Micropterus salmoides* | 97.0 | 3.84 | hunting macrofauna | benthopelagic | Centrarchidae | 2 |
| *Notemigonus crysoleucas* | 32.0 | 2.65 | variable | demersal | Leuciscidae | 32 |
| *Oncorhynchus clarkii* | 99.0 | 3.77 | hunting macrofauna | demersal | Salmonidae | 1 |
| *Oncorhynchus mykiss* | 122.0 | 4.08 | hunting macrofauna | benthopelagic | Salmonidae | 2 |
| *Oncorhynchus nerka* | 84.0 | 3.54 | selective plankton | pelagic-oceanic | Salmonidae | 1 |
| *Osmerus mordax* | 35.6 | 3.00 | hunting macrofauna | pelagic-oceanic | Osmeridae | 4 |
| *Perca flavescens* | 50.0 | 3.67 | hunting macrofauna | benthopelagic | Percidae | 14 |
| *Perca fluviatilis* | 60.0 | 4.35 | hunting macrofauna | demersal | Percidae | 1 |
| *Phoxinus phoxinus* | 14.0 | 3.20 | variable | demersal | Leuciscidae | 4 |
| *Pimephales notatus* | 11.0 | 2.66 | variable | demersal | Leuciscidae | 2 |
| *Pimephales promelas* | 10.1 | 2.42 | variable | demersal | Leuciscidae | 2 |
| *Platichthys flesus* | 60.0 | 3.32 | hunting macrofauna | demersal | Pleuronectidae | 1 |
| *Poecilia mexicana* | 11.0 | 2.00 | –* | benthopelagic | Poeciliidae | 1 |
| *Prosopium cylindraceum* | 59.0 | 3.30 | hunting macrofauna | benthopelagic | Salmonidae | 2 |
| *Rhinichthys atratulus* | 12.4 | 3.05 | variable | demersal | Leuciscidae | 6 |
| *Roeboides guatemalensis* | 13.0 | –* | –* | benthopelagic | Characidae | 1 |
| *Rutilus rutilus* | 50.2 | 2.96 | hunting macrofauna | benthopelagic | Leuciscidae | 1 |
| *Salmo salar* | 150.0 | 4.50 | hunting macrofauna | benthopelagic | Salmonidae | 1 |
| *Salvelinus fontinalis* | 86.0 | 3.31 | hunting macrofauna | benthopelagic | Salmonidae | 34 |
| *Salvelinus namaycush* | 150.0 | 4.29 | hunting macrofauna | benthopelagic | Salmonidae | 6 |
| *Semotilus atromaculatus* | 30.3 | 4.02 | hunting macrofauna | demersal | Leuciscidae | 22 |
| *Semotilus corporalis* | 51.0 | 3.41 | hunting macrofauna | demersal | Leuciscidae | 4 |
| *Squalius cephalus* | 60.0 | 2.73 | variable | benthopelagic | Leuciscidae | 3 |
| *Thymallus thymallus* | 60.0 | 3.14 | hunting macrofauna | benthopelagic | Salmonidae | 2 |
| *Umbra limi* | 14.0 | 3.67 | hunting macrofauna | demersal | Umbridae | 2 |

*Note.* Lmax = maximum total body length (cm) from FishBase. Trophic level from FoodTroph (or DietTroph where unavailable). DemersPelag = vertical habitat category from FishBase. Superscript asterisk (*) = value imputed; see Section 2.2. Feeding guild shortened from FishBase full labels. n netw. = number of corpus networks containing the species.

**Table S2.** Hyperparameters and architecture for all four models.

| Model | Architecture | Parameter | Value |
|---|---|---|---|
| Model A | Random Forest | n_estimators | 300 |
| | | max_features | sqrt |
| | | min_samples_leaf | 3 |
| | | Negative sampling | 1:5 (manual, per network) |
| | | Features | log mass ratio, movement match, metabolic match (3 features) |
| | | Training data | GATEWAy only (34 networks, LOSO) |
| Model B | Random Forest | n_estimators | 300 |
| | | max_features | sqrt |
| | | min_samples_leaf | 3 |
| | | Negative sampling | 1:5 (manual, per network) |
| | | Features | trophic diff, con/res Lmax, con/res trophic level, guild match, DemersPelag match, log Lmax ratio (8 features) |
| | | Training data | All 37 networks (LOSO) |
| Model C | Random Forest | n_estimators | 300 |
| | | max_features | sqrt |

Table S2 continued

| Model | Architecture | Parameter | Value |
|---|---|---|---|
| | | min_samples_leaf | 3 |
| | | Negative sampling | 1:5 (manual, per network) |
| | | Features | Model B features + habitat overlap score (9 features) |
| | | Training data | All 37 networks (LOSO) |
| Model D | Graph Attention Network | Layers | 2 (4 heads / 1 head) |
| | | Hidden dim. | 32 |
| | | Dropout | 0.3 |
| | | Optimizer | Adam (lr = 0.001, weight decay = 0.0001) |
| | | LR schedule | StepLR (step_size = 30, gamma = 0.5) |
| | | Epochs | 80 |
| | | Negative sampling | 1:5 (manual, per network) |
| | | Inference mode | Trait encoder only (inductive trait-only inference; no message passing) |
| | | Training data | All 37 networks (LOSO) |

*Note.* RF = random forest (scikit-learn RandomForestClassifier). GAT = graph attention network (PyTorch Geometric GATConv). All models used random seed 42. LOSO = leave-one-study-out cross-validation. Blank cells indicate the same value as the row above within each model block.

**Table S3.** Pairwise Wilcoxon signed-rank tests comparing model performance across networks.

| Comparison | Metric | n | Med. A | Med. B | Delta | W | p-value | Sig. |
|---|---|---|---|---|---|---|---|---|
| Model A vs. B | ROC-AUC | 34 | 0.973 | 0.609 | -0.364 | 0.0 | <0.001 | Yes |
| Model A vs. C | ROC-AUC | 34 | 0.973 | 0.606 | -0.367 | 0.0 | <0.001 | Yes |
| Model A vs. D | ROC-AUC | 34 | 0.973 | 0.553 | -0.420 | 0.0 | <0.001 | Yes |
| Model A vs. B | PR-AUC | 34 | 0.829 | 0.215 | -0.614 | 0.0 | <0.001 | Yes |
| Model A vs. C | PR-AUC | 34 | 0.829 | 0.213 | -0.616 | 0.0 | <0.001 | Yes |
| Model A vs. D | PR-AUC | 34 | 0.829 | 0.144 | -0.685 | 0.0 | <0.001 | Yes |
| Model B vs. C | ROC-AUC | 37 | 0.607 | 0.606 | -0.001 | 89.0 | 0.356 | No |
| Model B vs. D | ROC-AUC | 37 | 0.607 | 0.554 | -0.053 | 80.0 | <0.001 | Yes |
| Model C vs. D | ROC-AUC | 37 | 0.606 | 0.554 | -0.052 | 79.0 | <0.001 | Yes |
| Model B vs. C | PR-AUC | 37 | 0.207 | 0.206 | -0.001 | 68.0 | 0.167 | No |
| Model B vs. D | PR-AUC | 37 | 0.207 | 0.144 | -0.063 | 61.0 | <0.001 | Yes |
| Model C vs. D | PR-AUC | 37 | 0.206 | 0.144 | -0.062 | 62.0 | <0.001 | Yes |

*Note.* Med. = median AUC; W = Wilcoxon signed-rank test statistic; Sig. = significant at $p < 0.05$ after Holm-Bonferroni correction; Delta = median(B) - median(A). Model A comparisons restricted to GATEWAy networks (n = 34)

**Table S4.** Metadata for all 37 freshwater fish food webs included in the corpus.

| Network | Source | Ecosystem type | Nodes | Edges | Connectance | Fish spp. |
|---|---|---|---|---|---|---|
| Alford lake | gateway | lakes | 56 | 217.0 | 0.0705 | 2 |
| Balsam lake | gateway | lakes | 50 | 258.0 | 0.1053 | 2 |
| Beaver lake | gateway | lakes | 56 | 262.0 | 0.0851 | 4 |
| Big hope lake | gateway | lakes | 61 | 322.0 | 0.088 | 3 |
| Brandy lake | gateway | lakes | 30 | 119.0 | 0.1368 | 3 |
| Bridge brook lake | gateway | lakes | 75 | 548.0 | 0.0987 | 5 |
| Cascade lake | gateway | lakes | 35 | 115.0 | 0.0966 | 2 |
| Chub pond | gateway | lakes | 54 | 412.0 | 0.144 | 7 |
| Connera lake | gateway | lakes | 65 | 412.0 | 0.099 | 5 |
| Fawn lake | gateway | lakes | 32 | 119.0 | 0.12 | 2 |
| Grass lake | gateway | lakes | 44 | 162.0 | 0.0856 | 2 |
| Gull lake | gateway | lakes | 45 | 209.0 | 0.1056 | 2 |
| Helldiver pond | gateway | lakes | 41 | 167.0 | 0.1018 | 2 |
| Hoel lake | gateway | lakes | 72 | 564.0 | 0.1103 | 9 |
| Horseshoe Lake | gateway | lakes | 49 | 249.0 | 0.1059 | 5 |
| Little Rainbow Lake | gateway | lakes | 52 | 246.0 | 0.0928 | 2 |
| Long Lake | gateway | lakes | 65 | 413.0 | 0.0993 | 9 |
| Loon Lake | gateway | lakes | 35 | 115.0 | 0.0966 | 2 |
| Lost Lake | gateway | lakes | 31 | 145.0 | 0.1559 | 3 |
| Lower Sister Lake | gateway | lakes | 37 | 159.0 | 0.1194 | 2 |
| Rat Lake | gateway | lakes | 50 | 267.0 | 0.109 | 5 |
| Razorback Lake | gateway | lakes | 42 | 176.0 | 0.1022 | 3 |
| Safford Lake | gateway | lakes | 44 | 221.0 | 0.1168 | 4 |
| Sand Lake | gateway | lakes | 29 | 79.0 | 0.0973 | 2 |
| Squaw Lake | gateway | lakes | 41 | 166.0 | 0.1012 | 2 |
| Stink Lake | gateway | lakes | 53 | 277.0 | 0.1005 | 3 |
| Whipple Lake | gateway | lakes | 32 | 134.0 | 0.1351 | 5 |
| Bure Stream | gateway | streams | 95 | 1927.0 | 0.2158 | 2 |
| Dempsters Stream | gateway | streams | 110 | 965.0 | 0.0805 | 3 |
| Loddon Stream | gateway | streams | 105 | 2542.0 | 0.2328 | 5 |
| Lyde Stream | gateway | streams | 110 | 2030.0 | 0.1693 | 3 |
| Mill Stream | gateway | streams | 81 | 1435.0 | 0.2215 | 8 |
| Test Stream | gateway | streams | 121 | 3180.0 | 0.219 | 7 |
| Wensum Stream | gateway | streams | 127 | 3487.0 | 0.2179 | 7 |

Table S4 continued

| Network | Source | Ecosystem type | Nodes | Edges | Connectance | Fish spp. |
|---|---|---|---|---|---|---|
| Lake Crescent | mangal | freshwater | 6 | 7.0 | 0.2333 | 3 |
| Gatun Lake | mangal | freshwater | 17 | 32.0 | 0.1176 | 5 |
| Potreirinho Creek | mangal | freshwater | 106 | 235.0 | 0.0211 | 5 |

*Note.* Source: GATEWAy = Brose et al. (2019); Mangal = Poisot et al. (2016). Connectance = directed edges / n(n-1). Fish spp. = unique fish species with FishBase Lmax coverage.

**Table S5.** Model A and Model B performance by pair class across the 34 GATEWAy networks.

| Pair class | n networks | Median pairs | Median positive rate | Model A ROC-AUC | Model A PR-AUC | Model B ROC-AUC | Model B PR-AUC |
|---|---|---|---|---|---|---|---|
| All pairs | 34 | 2551 | 0.105 | 0.973 | 0.830 | 0.609 | 0.215 |
| Fish-associated | 34 | 319 | 0.190 | 0.912 | 0.705 | 0.887 | 0.529 |
| Non-fish | 34 | 2209 | 0.092 | 0.979 | 0.868 | 0.500 | 0.092 |

*Note.* Values are medians across the 34 GATEWAy folds. Model A uses local GATEWAy body mass and Model B uses FishBase traits; both were computed from the same full, unbalanced held-out predictions.

**Table S6.** Permutation importance for Models B and C (random forest), ranked by Model B importance.

| Feature | Model B (Δ ROC-AUC) | Model C (Δ ROC-AUC) | Description |
|---|---|---|---|
| Resource trophic level | 0.0104 | 0.0069 | Resource trophic level (FoodTroph) |
| Trophic level difference | 0.0071 | 0.0094 | Consumer minus resource trophic level |
| DemersPelag match | 0.0040 | 0.0018 | Consumer and resource share depth stratum |
| Consumer Lmax (cm) | 0.0040 | 0.0038 | Max body length of consumer (FishBase) |
| Feeding guild match | 0.0028 | 0.0056 | Consumer and resource share feeding guild |
| Resource Lmax (cm) | 0.0003 | 0.0007 | Max body length of resource (FishBase) |
| Log Lmax ratio | -0.0001 | 0.0002 | ln(consumer Lmax / resource Lmax) |
| Consumer trophic level | -0.0018 | -0.0006 | Consumer trophic level (FoodTroph) |
| Habitat overlap score | n/a | 0.0006 | Depth-stratum overlap score, 0–1 (Model C only) |

*Note.* Δ ROC-AUC = mean decrease in ROC-AUC per leave-one-study-out fold (n = 37); negative values indicate permutation did not reduce performance on average. Ranked by Model B importance.
